# A Genomic Language Model for Zero-Shot Prediction of Promoter Variant Effects

**DOI:** 10.1101/2024.11.11.623015

**Authors:** Courtney A. Shearer, Rose Orenbuch, Felix Teufel, Christian J. Steinmetz, Daniel Ritter, Erik Xie, Artem Gazizov, Aviv Spinner, Jonathan Frazer, Mafalda Dias, Pascal Notin, Debora S. Marks

## Abstract

Disease-associated genetic variants occur extensively in noncoding regions like promoters, but current methods focus primarily on single nucleotide variants (SNVs) that typically have small regulatory effect sizes. Expanding beyond single nucleotide events is essential with insertions and deletions (indels) representing the logical next step as they are readily identifiable in population data and more likely to disrupt regulatory elements. However, existing methods struggle with indel prediction, and clinical interpretation often requires assessing complete promoter haplotypes rather than individual variants. We present LOL-EVE (Language Of Life for Evolutionary Variant Effects), a conditional autoregressive transformer trained on 13.6 million mammalian promoter sequences that enables both zero-shot indel prediction and complete promoter sequence scoring. We introduce three benchmarks for promoter indel prediction: ultra rare variant prioritization, causal eQTL identification, and transcription factor binding site disruption analysis. LOL-EVE’s superior performance demonstrates that evolutionary patterns learned from indels enable accurate assessment of broader promoter function. Application to Genomics England clinical data shows that LOL-EVE can prioritize promoter haplotypes in known developmental disorder genes, suggesting potential utility for clinical variant assessment. LOL-EVE bridges individual variant prediction with haplotype-level analysis, demonstrating how evolution-based genomic language models may assist in evaluating regulatory variants in complex genetic cases.

## 1 Introduction

The molecular language of life, DNA, has existed for over 4 billion years, constantly subject to evolutionary pressures. Evolution through natural selection can be seen as a series of countless experiments refining the genomic code to maximize organismal fitness. A long-standing challenge of computational biology is how to use genomic information to learn a mapping between genomic state and the corresponding organism state, i.e. genotype to phenotype. Using evolutionary sequences for phenotype predictions allows assessment of mutational impacts on organism fitness without requiring a priori knowledge of impact mechanisms or experimental work. While substantial progress has been made in developing computational methods to determine protein variant effects on phenotype [17, 26, 51, 58, 50, 49], methods for predicting the effects of variants in non-coding regions are still in their infancy.

Non-coding regions, which make up 99% of the genome, contain thousands of variants linked to human disease [44]. These variants contribute to many rare and undiagnosed diseases that have eluded diagnosis through exome sequencing alone [43]. However, identifying whether these non-coding variants cause phenotype changes or are in linkage disequilibrium with causal variants remains challenging [1].

In clinical practice, variant interpretation often involves assessing multiple variants across promoter regions rather than isolated single nucleotide changes. Traditional approaches that score variants independently fail to capture the cumulative regulatory impact of complex haplotypes, limiting their utility for rare disease diagnosis where patients may carry combinations of variants that collectively disrupt gene regulation.

Current approaches to variant effect prediction in non-coding regions primarily examine single nucleotide variants (SNVs), largely due to the relative ease of their detection in whole-genome sequencing [45, 29]. While this SNV-focused approach has yielded valuable insights, several studies suggest that individual SNVs are unlikely to have large effects at an organismal scale, especially in non-coding regions [34, 56], due to the redundancy built into biological systems and the generally smaller effect sizes of non-coding variants [68]. However, there is considerable heritability in promoter regions, more than would be expected from individual SNVs—indels are a likely contributor [20, 14]. Insertions and deletions represent an important but understudied source of genetic variation [37], and are more likely to disrupt regulatory elements than individual SNVs due to their ability to affect multiple nucleotides simultaneously. While genomic language models have advanced rapidly [7, 47, 13], most focus on SNVs and lack specialized training for promoter indel prediction or the capability to assess complete promoter sequences.

Promoter variation accounts for a significant percentage of undiscovered diseases [44, 2], although research to date has revealed only small effects on clinical outcomes and gene expression [18, 24]. Recent research has shown that the orientation and order of transcription factor (TF) binding sites are major drivers of gene regulatory activity [22]. SNPs rarely cause changes disrupting these patterns, thus necessitating a method that can predict the effects of multiple nucleotide changes.

Furthermore, many methods have relied on expression or chromatin accessibility data, which, while highly informative in specific biological contexts [57], are often difficult to gather for diverse variant types and experimental conditions. This limitation is particularly pronounced for indels, where functional assays are more challenging than for single nucleotide variants. Assessments of existing supervised approaches like CADD [55] show that they suffer from data leakage issues that inflate performance [23], undermining their reliability for true zero-shot prediction in cases not represented in the training data. These challenges motivate the development of zero-shot evolutionary methods that can generalize to unseen variants without requiring additional experimental data, providing tremendous practical value for variant interpretation.

We hypothesize that expanding the scope of variant effect prediction to include indels, particularly in promoter regions, will lead to the discovery of variants with larger phenotypic effects [65, 11]. This approach will potentially identify previously overlooked sources of genetic variation with significant phenotypic impacts, contributing to a deeper understanding of rare and undiagnosed diseases and uncovering new pathways for diagnosis and treatment. Moreover, the autoregressive nature of our approach enables scoring of complete promoter haplotypes containing multiple variants—a critical capability for clinical variant interpretation in rare diseases.

In this work, we present LOL-EVE (Language Of Life for Evolutionary Variant Effects), a genomic language model for zero-shot prediction of promoter indel effects and complete promoter sequence scoring. LOL-EVE bridges research-focused variant prediction with potential clinical applications by enabling assessment of both individual variants and complex haplotypes. Our key contributions are as follows:

- We construct and open source, PromoterZoo, a dataset of **13.6 million promoter sequences** comprising almost 20 thousand 1kb promoter region sequences from 447 species across mammalian evolution identified in the Zoonomia project [12] (§ 2.1);
- We develop LOL-EVE, **a 235 million parameter conditional generative model of promoter evolution** for predicting variant effects (§ 2.2);
- We introduce **three new benchmarks** specifically designed for zero-shot indel variant effect prediction in promoter regions, encompassing ultra rare indel detection, causal variant prioritization and TF binding site disruption (§ 3).
- We evaluate LOL-EVE’s **clinical utility** by demonstrating effective prioritization of promoter haplotypes in known developmental disorder genes using real patient data (§ 4.3).

## 2 LOL-EVE

### 2.1 Training data

Promoters and other regulatory regions generally evolve faster than protein-coding sequences, as regulatory changes can often be more easily tolerated than changes to protein structure and function [63]. To capture these evolutionarily relevant regulatory signals, particularly those that have evolved recently, we focused on training data from mammals. We curated a promoter dataset across 447 diverse species from the Zoonomia project [12, 35].

Since TSS annotations are not readily available for most species in our dataset, we employed a comparative genomics approach to identify putative promoter regions. We leveraged sequence similarity to the first exon of 19,254 protein-coding genes from the NCBI RefSeq human genome annotation, using the HAL toolkit[25] to perform liftover of these exon coordinates to each species. We then extracted the 1,000 base pairs upstream of each exon start as putative promoter regions, accounting for strand orientation and avoiding overlap with neighboring gene bodies (Figure 1A-left).

**Figure 1:**
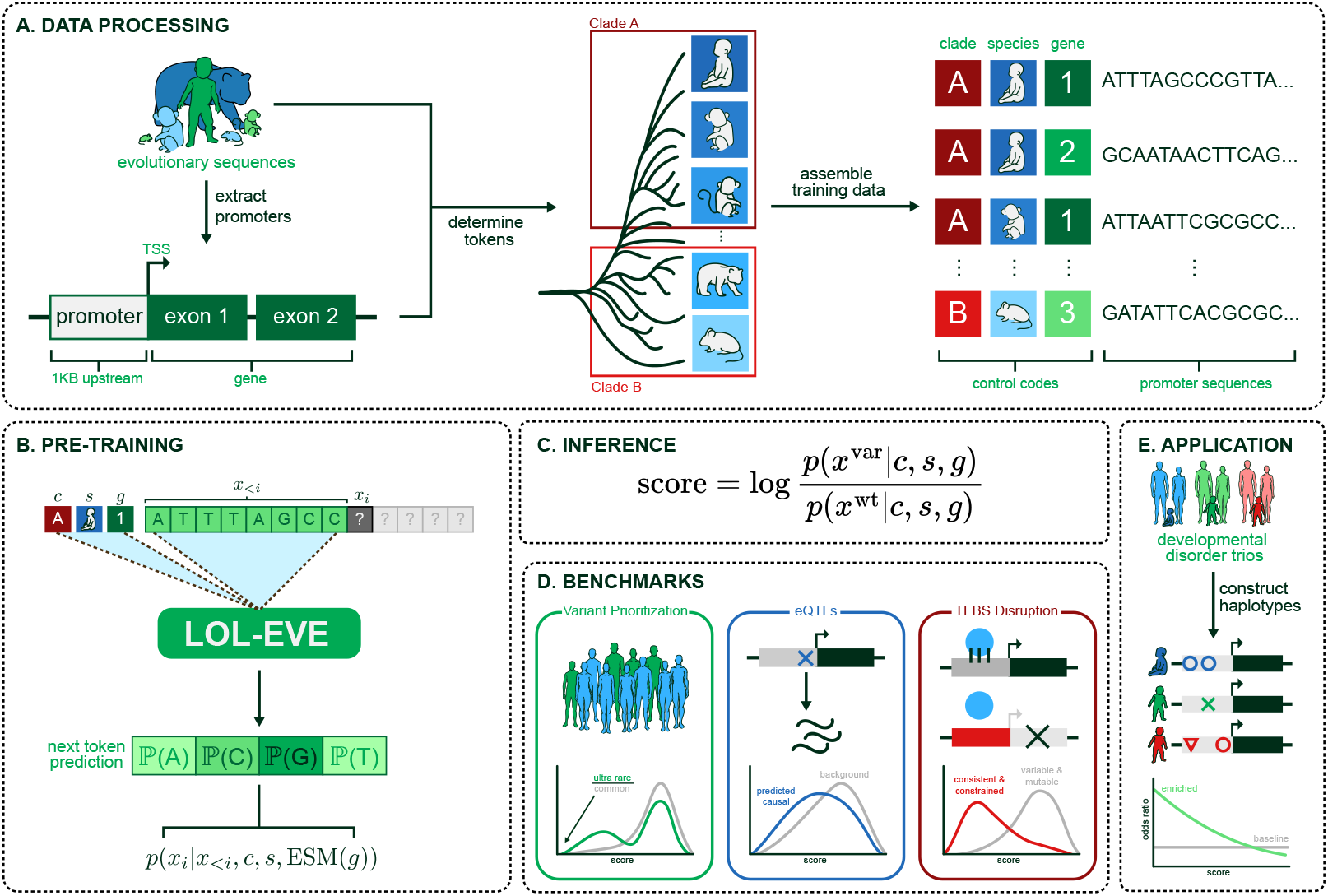
LOL-EVE approach overview. A. Data preprocessing: Promoter sequences extracted from evolutionary sequences across mammals, grouped into clades and tokenized with control codes. B. Pre-training: Next-token prediction conditioned on sequence context and control codes. C. Inference Equation D. Benchmarks: Evaluation on variant prioritization, eQTLs, and TFBS disruption tasks. E. Overview of Application to Development Disorder Trios

To validate our approach, we scored all extracted sequences using the Sei promoter score [8], which is trained on functional genomics data from humans. Despite being human-based, the promoter scores generalize well across species (Figure A1), showing strong conservation of regulatory elements in mammalian species. Including reverse complements, this resulted in a dataset of 13.6 million sequences. We employed a chromosome-wise split with chromosome 19 used for validation, ensuring no gene information leakage between training and validation sets (Sec A.2).

### 2.2 Model Architecture

To address the challenge of modeling non-aligned promoter sequences across mammalian evolution for indel variant effect prediction, LOL-EVE learns a generative model over full promoter nucleotide sequences. To incorporate evolutionary context, the model conditions its predictions on the promoter’s most proximal gene, species, and clade, such as non-primate mammals and primates (Figure 1A-right). This strategy is implemented using a decoder-only transformer architecture, following the CTRL framework [33] (Figure 1B). The conditioning information is provided as prefix tokens, allowing LOL-EVE to autoregressively generate and score promoter sequences in a context-aware manner. This approach enables the model to capture both broad evolutionary patterns and species-specific variations in regulatory elements. This clade specificity, as shown in (Figure 1A-mid), can be useful for capturing, in this model, mammal vs. primate-specific constraint, which has been shown to be crucial for distinguishing disease-associated regulatory variants. Specifically, *primate-constrained elements* are more likely to harbor regulatory variants tied to human-specific traits and diseases, while *mammal-constrained elements* may underlie conserved regulatory processes across a broader evolutionary scope [35].

To better capture the distinct roles of control codes and genomic sequences, we developed an adaptive local position embedding scheme defined as:

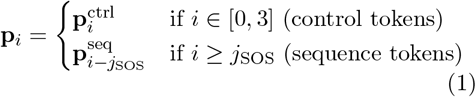

where 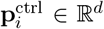 are absolute position embed-dings for control tokens and 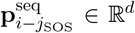 are relative position embeddings that reset at the sequence start token position *j*_SOS_. This adaptive approach allows the model to maintain structural understanding of control codes while enabling biologically meaningful positional representations for genomic sequences.

We provide the list of all model hyperparameters used in our final architecture in Table A1. Unlike LMs that use k-mer tokenization schemes to achieve length compression [13, 67], LOL-EVE directly tokenizes the promoter sequence *x* at base pair resolution. This enables the model to accurately handle insertions and deletions without causing tokenization shifts downstream.

To encode the most proximal gene *g*, we use mean-pooled ESM2 embeddings (`ESM2_t33_650M_UR50D`) [39] of a gene’s canonical human protein sequence. These embeddings are kept frozen during training and are projected from dimension 1280 to LOL-EVE’s embedding dimension using a learned linear mapping providing a tensor denoted at ESM(*g*). The ESM-based embedding scheme allows LOL-EVE to generalize to gene tokens unseen during training, which is critical in genomics where chromosome-wise hold outs are typically preferred. The species *s* and clade *c* are encoded using learned embeddings. Taken together, LOL-EVE (*p*_*θ*_) models the conditional distribution of a length *L* promoter autoregressively

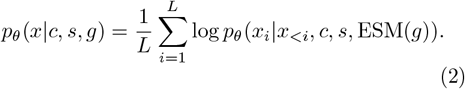

We apply control tag dropout with probability *α* to data set *D* to encourage the model to learn representations robust to the presence of such tags and mitigate memorization. During training the loss − with dropout is given by

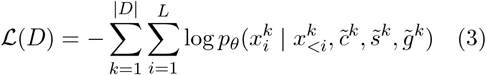

where 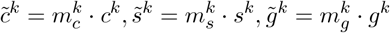and 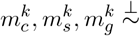 Bernoulli(*α*)

We implement a strand-aware length dropout mechanism to account for the inherent directionality of DNA sequences, as shown in Equation 4. For sequences on the forward strand (*d* = 1), tokens are shifted leftward after dropping out l tokens from the right end, maintaining causal attention over the remaining sequence. For reverse strand sequences (*d* = *−*1), tokens are simply dropped from the right end without shifting, preserving the natural 5’ to 3’ processing order. In both cases, dropped tokens are replaced with padding tokens that are ignored in self-attention layers, and the maximum dropout length is capped at 90% of the sequence length to ensure sufficient context is retained.

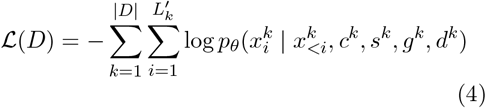

where *l*^*k*^ *∼* Uniform(0, 0.9|*x*^*k*^|),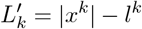 and *d*^*k*^ *∈ {−*1, 1*}* indicates strand direction

At inference we use the score in Equation 5 which represents the log-likelihood ratio between variant, *x*^var^, and wildtype, *x*^wt^, sequences. This captures how likely the variant sequence is compared to wildtype [17, 51, 48, 10, 59].

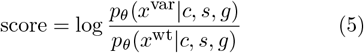

## 3 Indel Benchmarks

Our goal is to assess promoter variant effects through the lens of regulatory function and evolutionary constraint. To that end, we introduce three indel benchmarks that probe complementary dimensions of variant impact:

- **Ultra-rare variant prioritization** evaluates selection pressure in the human population – as deleterious variants are purified out of the population, they will be incredibly rare reflecting evolutionary intolerance.
- **Causal eQTL prioritization** focuses on expression-altering function – whether a variant is likely to be causal for changes in gene expression, based on fine-mapped expression quantitative trait loci.
- **TFBS disruption** assesses the functional integrity of transcriptional regulation – whether a variant disrupts transcription factor binding sites, particularly in genes where such disruption is expected to be deleterious due to high constraint and consistent expression.

Together, these tasks form a biologically grounded benchmark suite that reflects the core goals of variant effect prediction: identifying variants that either perturb gene regulation or are under negative selection in humans. Details on scoring methodologies are provided in subsection C.1.

### 3.1 Ultra Rare Variant Prioritization

#### Rationale

Ultra rare variants are more likely to be functionally important or disease-causing compared to more common variants [41]. As such, models should assign their most extreme predictions to only the rarest of variants.

#### Task

We evaluate how strongly models prioritize ultra rare variants (MAF *<* 0.00001) versus common variants (MAF *>* 0.001) by comparing their scores at each percentile cutoff. Specifically, for each percentile we take the ratio

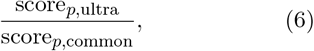

(*>* 1 means stronger ultra-rare signal), and we report the mean of these ratios stratified by indel-length

#### Data

We evaluate variants from gnomAD V4.0 [9], comparing predictions between ultra rare variants (MAF *<* 0.00001)[61] and more common variants (MAF *>* 0.001). By examining ratios across multiple percentile thresholds, we assess how consistently models prioritize ultra rare variants at different levels of stringency.

### 3.2 Causal eQTL Prioritization

#### Rationale

An expression quantitative trait locus (eQTL) is a variant associated with a change in gene expression. Fine-mapping methods such as SuSiE [62] assign each indel a posterior inclusion probability (PIP) reflecting its likelihood of being causal. Here, we focus on *cis*-eQTLs—indels within promoter regions whose eGene (linked gene) is proximal.

#### Task

Given two sets of promoter indels—putatively causal (*PIP >* 0.99) and background (*PIP <* 0.01)—models should assign larger absolute effect scores to the causal group. We assess discrimination by AUROC and AUPRC normalized by the causal-variant fraction.

#### Data

We retrieved fine-mapped cis-eQTL indels from the eQTL Catalogue [32], filtered to promoters whose eGene matches the variant’s nearest gene. Applying PIP thresholds of 0.99 and 0.01. We also stratify by the likelihood of slippage – repeated regions likely to have repeat expansion or contraction. For the cumulative-slippage analysis (see Section C.5 and Figure A3), we compute running-mean AU-ROC and normalized AUPRC at slippage cutoffs of 25 bp, 50 bp, 100 bp, 200 bp, and *>* 200 bp (Table A4).

### 3.3 TFBS Disruption

#### Rationale

Transcription factors (TFs) are essential regulators of gene expression, binding to specific DNA sequences in promoter regions to control transcriptional activity. Disruptions to TF binding sites (TFBS) can impact gene regulation, with the severity depending on the evolutionary constraint and expression characteristics of the target gene. We hypothesize that variants disrupting TFBS should be most deleterious in genes that are evolutionarily constrained and consistently expressed across tissues, as these genes are typically intolerant to regulatory perturbations [64].

#### Task

We evaluate whether models correctly predict that TFBS disruptions are more deleterious in genes with high evolutionary constraint and low expression variability compared to genes with low constraint and high variability. For each transcription factor, we compute model disruption scores for TFBS variants in each gene class and assess whether high-constraint/low-variability genes receive consistently lower (more deleterious) scores than low-constraint/high-variability genes.

Performance is measured as delta accuracy across transcription factors using balanced sampling with the following setup:

Let ℋ be the set of *high-constraint/low-variability* genes and ℒ the set of *low-constraint/high-variability* genes (see Sec. C.4). For each transcription factor *t* = 1, …, *T*, let Score_*t*_(𝒢) be the model’s mean disruption score across genes in set 𝒢. We define

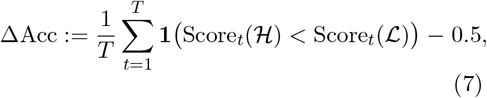

where **1**(·) is the indicator function. A positive ΔAcc means the model assigns lower disruption scores to the high-constraint set ℋ more often than expected by chance.

#### Data

Genes are categorized using mammalian evolutionary rates (OrthoDB) and expression variability (GTEx CV). TFBS disruptions are identified using JASPAR CORE TFs and position-specific scoring matrices. Complete methodology and data processing details are in Appendix C.4.

## 4 Results

### 4.1 Indel Benchmark

We benchmark LOL-EVE against a diverse set of models: DNA language models applicable to the human genome (HyenaDNA [47], DNABERT-2 [67], Nucleotide Transformer (NT) [13], Caduceus [54], and Genomic Pretrained Network (GPN) [5, 6]), SpeciesLM [19], and Evo1 [46] and Evo2[7], supervised predictors (CADD [34] and Enformer [3]), and conservation metrics (PhyloP). While some supervised models (CADD, Enformer) achieve competitive performance, this performance is likely inflated due to data leakage issues where their training data overlaps with the datasets used to define our evaluation benchmarks (see Section C.1.2). Therefore, we focus our main discussion on unsupervised approaches that operate in a true zero-shot capacity. For LMs that make multiple checkpoints available, we focus our discussion on the best performing checkpoint in each experiment, with remaining checkpoints evaluated in section B.6.

#### 4.1.1 Ultra Rare Variant Prioritization

Among unsupervised models, LOL-EVE delivers superior performance overall, achieving the highest enrichment for medium indels (2.150 *±* 0.051) and large indels (1.956 *±* 0.118), and ranks competitively small (1.482 *±* 0.032). GPN-Promoter, a masked-language transformer trained specifically on promoter sequences, excels on small indels (2.297 *±* 0.051) where its local-context objective is most effective. HyenaDNA shows consistent moderate performance across all categories (1.323 *−* 1.400), while other LMs like NT and Caduceus achieve only modest ratios (≈ 1.02–1.26). DNABERT-2 and speciesLM remain near baseline, and notably, Evo2 performs below baseline for small and medium indels (0.822 and 0.757 respectively).

LOL-EVE’s exceptional performance on medium-length indels among unsupervised models suggests the model has learned to recognize biologically meaningful patterns at the scale most relevant for regulatory disruption. This size range encompasses typical transcription factor binding motifs and regulatory elements, where insertions or deletions are likely to disrupt critical protein-DNA interactions. Furthermore, its performance across all indel sizes indicates it has captured evolutionary constraints that operate at multiple scales—from single nucleotide changes affecting individual binding sites to larger structural disruptions affecting multiple regulatory elements.

The clear advantage of promoter-specialized models (LOL-EVE and GPN-Promoter) over general genomic models highlights the importance of region-specific training. While foundation models like Evo2 achieve impressive performance on genome-wide tasks, their below-baseline performance on promoter indels demonstrates that general models may not capture the specific evolutionary constraints operating in regulatory regions.

Notably, while supervised models like CADD show strong performance in the appendix results, they rely on the very population frequency data that defines our benchmark categories (ultra-rare vs. common variants), creating a form of data leakage that inflates their apparent performance. LOL-EVE’s purely evolutionary approach circumvents this issue, providing genuine zero-shot predictions based solely on patterns learned sequence conservation.

### 4.2 Causal eQTL Prioritization

Figure 2 shows cumulative ROC AUC and normalized AUPRC for causal versus background cis-eQTL indels as we include variants within increasing slippage cutoffs (C.5). LOL-EVE leads among all unsupervised models at every threshold—peaking near 0.73 ROC AUC and 3.1 × baseline AUPRC at 25 bp—and sustains the strongest separation even at large distances. Among other unsupervised approaches, SpeciesLM and GPN-Promoter shows modest but consistent performance, while other LMs (e.g., NT-2.5B, DNABERT-2) achieve more limited gains. Notably, LOL-EVE also generalizes to SNPs (Figure A5), though overall performance across all models is modest for single nucleotide variants, consistent with our hypothesis that individual SNPs have smaller regulatory effect sizes compared to indels.

**Figure 2:**
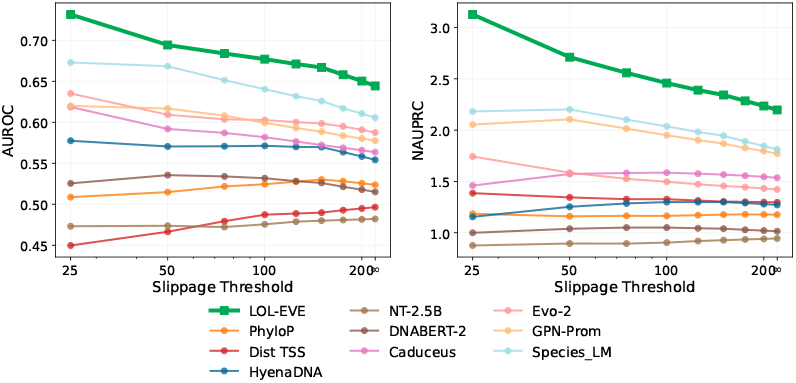
Cumulative causal-eQTL prioritization per-formance across slippage thresholds (log scale). Left: running mean ROC AUC; Right: running mean normalized AUPRC (AUPRC / baseline). Full models in Figure A4, Variants/Genes per threshold are shown in in Table Table A4

LOL-EVE’s consistent performance across all slippage thresholds among unsupervised models provides compelling evidence that the model has learned biologically relevant mutational mechanisms rather than merely statistical correlations. In other words, LOL-EVE’s predictions correlate with known mechanisms of indel formation—particularly in repetitive sequences and homopolymer runs where DNA polymerase slippage commonly occurs during replication. Its sustained performance at high slippage scores (≤ 200*bp* total repeat length) distinguishes it from conservation-based approaches like PhyloP, which rely on position-specific conservation scores and may struggle in repetitive regions where alignment-based methods face challenges. While supervised models (CADD and Enformer) show competitive performance in appendix results, they benefit from training on expression and regulatory datasets that directly relate to the eQTL task being evaluated. This creates a form of task-specific data leakage that artificially inflates their performance. LOL-EVE’s success using only evolutionary sequence information demonstrates the power of cross-species patterns for identifying functionally relevant variants without task-specific supervision.

**Table 1:** Mean ratio and standard error across indel length categories and percentiles (1%, 2.5%, 5%, 10%). Best checkpoints: `tiny` (HyenaDNA), `ph-131k` (Caduceus), `2.5B-mulit` (NT). See Table A3 for all models and Table A4 for variant/gene counts per threshold.

| Model | Small (1–2bp) |  | Medium (3–15bp) |  | Large (16–50bp) |  |
| --- | --- | --- | --- | --- | --- | --- |
|  | Mean | SE | Mean | SE | Mean | SE |
| LOL-EVE | 1.482 | 0.032 | **2.150** | 0.051 | **1.956** | 0.118 |
| GPN-Promoter | **2.297** | 0.051 | 1.912 | 0.031 | 1.456 | 0.072 |
| SpeciesLM | 1.220 | 0.017 | 0.993 | 0.078 | 1.124 | 0.051 |
| DNABERT-2 | 1.069 | 0.012 | 0.974 | 0.017 | 1.050 | 0.004 |
| Caduceus | 1.028 | 0.045 | 1.116 | 0.074 | 1.253 | 0.172 |
| NT | 1.022 | 0.014 | 1.053 | 0.014 | 1.261 | 0.026 |
| HyenaDNA | 1.323 | 0.028 | 1.400 | 0.015 | 1.361 | 0.014 |
| Evo2 | 0.822 | 0.041 | 0.757 | 0.014 | 1.043 | 0.039 |

#### 4.2.1 TFBS Disruption

Figure 3 shows that LOL-EVE most accurately distinguishes TFBS disruptions in high-constraint, consistently expressed genes from those in low-constraint, variably expressed genes. For the greatest proportion of TFs, LOL-EVE correctly assigns lower scores to the high-constraint set, reflecting their greater sensitivity to loss of binding sites. This aligns with the expectation that variably expressed genes tolerate TFBS disruptions more readily than consistently expressed ones. By capturing these differential sensitivities, LOL-EVE demonstrates predictive power for promoter variant impact, indicating it has internalized the relationship between evolutionary constraint and functional importance. Interestingly, the two other models showing modest positive performance—HyenaDNA and PhyloP—each capture complementary aspects that LOL-EVE combines. HyenaDNA’s autoregressive architecture enables sequential modeling of regulatory context, while PhyloP provides deep evolutionary conservation signals. Notably, Enformer performs below baseline on this task and, as shown in Figure A6, frequently predicts effects in the wrong direction—a trend consistent with prior findings that sequence-to-expression models like Enformer struggle with predicting directionality of variant effects across individuals [53, 27].

**Figure 3:**
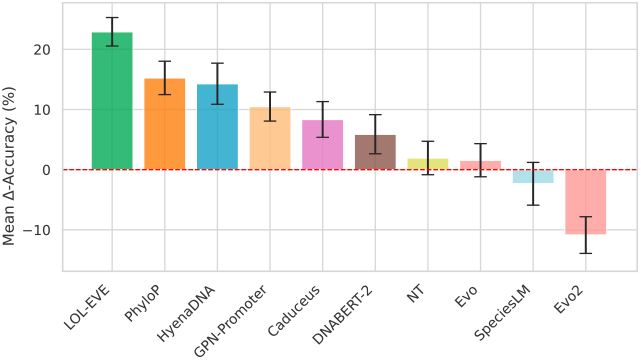
Mean delta-accuracy in TFBS disruption (±*SE*) for high-constraint/low-variability versus low-constraint/high-variability genes. Full results in Figure A6, with gene counts per threshold in Figure A7.

#### 4.2.2 Synthesis Across Benchmarks

Collectively, these results demonstrate that LOL-EVE has learned an understanding of promoter evolution that operates across multiple biological scales. The model’s success spans from identifying variants under negative selection (ultra-rare prioritization) to predicting functional regulatory effects (eQTLs) to making context-dependent assessments of binding site importance (TFBS disruption).

Ablation analysis in Figure A8 reveals that different prediction tasks benefit from different aspects of evolutionary conditioning. Evolutionary context of species and clade is beneficial for tasks concerned with constraint (ultra rare variant prioritization), while gene tokens are most beneficial for gene-specific function (eQTLs). Tasks that require both understanding of constraint and gene-specific function (TFBS disruption) benefit from the full evolutionary context: gene, species and clade.

This adaptive use of evolutionary information suggests that LOL-EVE has developed specialized representations for different types of biological questions, rather than learning a single global model of sequence constraint. Having established LOL-EVE’s superior performance on individual indel prediction, we next evaluate whether these learned evolutionary patterns can be applied to assess complete promoter sequences in a clinical context.

### 4.3 Application to Clinical Cohort

Clinical variant interpretation often requires evaluating multiple variants within promoter regions rather than individual variants in isolation. To evaluate LOL-EVE’s ability to score complete haplotypes, we utilized promoter haplotype data from severe developmental disorder patients in the Genomics England 100,000 Genomes dataset [21]. We constructed haplotypes by phasing variants according to parental inheritance patterns within promoter regions and scored complete sequences using LOL-EVE’s autoregressive architecture.

Promoter haplotypes were stratified based on whether they occurred upstream of genes with known associations to developmental disorders according to the Developmental Disorders Gene2Phenotype (DDG2P) database [15]. For a range of score thresholds, we calculated the enrichment of deleterious scores for promoters of developmental disorder genes compared to the background. Then, we compared performance against other autoregressive models (Evo2 and HyenaDNA). Detailed methodology is provided in Appendix D.

#### 4.3.1 Enrichment in Developmental Disorder Genes

Figure 4 shows odds ratios for deleterious haplotypes in DDG2P genes versus background genes across different log-likelihood ratio thresholds. LOL-EVE demonstrates consistent enrichment for DDG2P genes, with odds ratios reaching approximately 4-fold at stringent thresholds, indicating that the most deleterious scoring haplotypes are significantly more likely to occur upstream of known developmental disorder genes. In comparison, HyenaDNA consistently shows odds ratios below 1, indicating no enrichment, while Evo2 demonstrates modest performance at lower thresholds.

**Figure 4:**
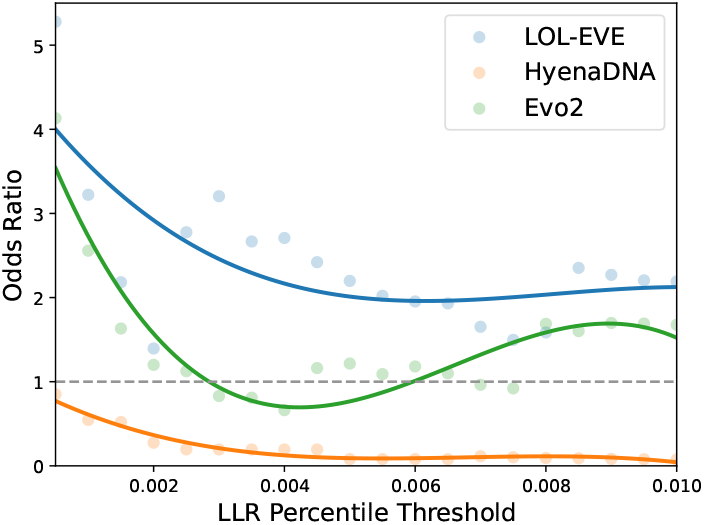
The Odds Ratio plotted against the change in LLR percentile threshold for each of LOL-EVE, HyenaDNA, and Evo2.

Figure A9 shows strong correlation between patient counts and gene counts across models. Figure A10 demonstrates that LOL-EVE and Evo2 maintain high gene detection even at stringent thresholds, while HyenaDNA shows minimal detection throughout the analysis.

## 5 Conclusion

LOL-EVE demonstrates superior performance for promoter indel effect prediction by leveraging evolutionary patterns learned from mammalian sequences. The model’s success spans from identifying variants under negative selection to predicting functional regulatory effects, with particularly strong performance on medium-length indels that are most likely to disrupt transcription factor binding sites.

Beyond individual variant assessment, LOL-EVE’s autoregressive architecture enables scoring of complete promoter sequences—a capability demonstrated through application to Genomics England clinical data. The enrichment of deleterious promoters scores in front of known developmental disorder genes provides preliminary evidence that evolutionary patterns learned from LOL-EVE can inform clinical variant prioritization.

As precision medicine increasingly relies on interpreting non-coding variants, evolution-based approaches like LOL-EVE offer genuine zero-shot predictions while avoiding the data leakage issues of supervised methods. This work demonstrates the potential of genomic language models trained on evolutionary data for identifying regulatory variants in complex genetic cases.

## 6 Data and Code Availability

Code for this work is publicly available at:

- GitHub:https://github.com/debbiemarkslab/LOL-EVE
- HuggingFace:https://huggingface.co/Marks-lab/LOL-EVE

## 7 Acknowledgement

This research was made possible through the generous support and collaboration of several individuals and institutions. The authors thank members of the Marks lab, particularly Sarah Gurev, and the Medical and Population Genetics Group at the Broad Institute for their valuable discussions and constructive feedback throughout this project. The computational resources provided by the Massachusetts Green High Performance Computing Center (MGHPCC) and Novo Nordisk are gratefully acknowledged as essential for completing this work. Additionally, this research was made possible through access to data in the National Genomic Research Library, which is managed by Genomics England Limited (a wholly owned company of the Department of Health and Social Care). The National Genomic Research Library holds data provided by patients and collected by the NHS as part of their care and data collected as part of their participation in research. The National Genomic Research Library is funded by the National Institute for Health Research and NHS England. The Wellcome Trust, Cancer Research UK and the Medical Research Council have also funded research infrastructure

## Appendix

### A Model details

#### A.1 Hyperparameters

**Table A1:** The hyperparameters of the LOL-EVE model.

| Hyperparameter | Value |
| --- | --- |
| Dimension | 768 |
| Layers | 12 |
| Heads | 12 |
| Feedforward dimension | 8192 |
| Learning rate | $1e^{-5}$ |
| Batch size | 32 |
| Steps | 150,00 |

#### A.2 Training Data

##### A.2.1 Comparative Genomics Approach

Transcription Start Site (TSS) annotations, which are often used to infer promoter regions, are not readily available for most species in our dataset due to several factors. Many of the 447 species lack comprehensive genome annotations, particularly for regulatory regions like promoters. Even in well-annotated species, TSS and promoter definitions can vary significantly across different databases and research groups.

To address this, we employed a comparative genomics approach to identify putative promoter regions, leveraging sequence similarity to the first exon of 19,254 protein-coding genes from the NCBI RefSeq human genome annotation (assembly GRCh38.p14, annotation release 109). This strategy allowed us to consistently infer promoter regions across species by aligning known human exonic regions to homologous exons in other species, then extracting sequences upstream of the start of the first exon (which we define as the putative TSS). It’s important to note that no genome has “promoter annotations” as such; rather, we use these inferred TSS positions and their upstream sequences as proxies for promoter regions. Importantly, in the human annotations we utilized, the 5’UTR often overlaps with the annotations for exon 1, which influences our definition of putative promoter regions across species. A visual representation of the sequence regions is shown in Figure 1A-left.

##### A.2.2 Sequence Extraction and Processing

Using the HAL toolkit [25], we performed a liftover of these exon coordinates to each species in the Zoonomia project. For each species, exons were retained if their length was at least 50% of the length of the corresponding human exon. This threshold ensured that conserved regions were captured while excluding regions where the alignment is unreliable.

To define promoter regions, we extracted the 1,000 base pairs upstream of each exon start, accounting for the strand orientation of the gene. If the upstream region overlapped with the neighboring gene body, we shortened the promoter region to avoid misclassifying coding regions or intergenic space as promoters. This conservative approach minimized the risk of including non-promoter sequences but may exclude more distal regulatory elements, a potential caveat of the 1,000 bp window approach. Additionally, in cases where promoter regions from neighboring genes were within 100 base pairs of each other, we merged the coordinates. This merging process ensured that promoter regions were not artificially fragmented due to closely spaced genes.

##### A.2.3 Validation Using Sei Scores

To gain further insight into the validity of the upstream 1,000 bp approach, we scored all extracted sequences using the Sei promoter score [8], which is trained on functional genomics data from humans. Despite Sei being human-based, we found that the promoter scores generalize well across species, showing strong conservation of regulatory elements in many mammalian species. Notably, promoters from species closely related to humans, such as other primates, tend to have higher Sei scores, indicating similar promoter activity, while more distant species still retain significant functional signal, suggesting that core regulatory sequences are preserved across mammals (**??**). Further we assessed how the Sei score distributions for 3 groups: Human Coding Sequence (CDS) regions, Human promoters, and our training data compare (**??**). Our training data promoter distribution aligns more closely with the raw Human promoters than the Human CDS regions, providing additional validation of our comparative genomics approach.

##### A.2.4 Data Splitting

Including reverse complements, this resulted in a dataset of 13.6 million sequences. We employed a chromosome-wise split for development, with chromosome 19 used for validation. Promoters from non-human species were assigned to the respective set based on the chromosome of the human gene used for liftover, thereby ensuring that all instances of a gene are placed in the same partition and no gene information leakage between the training and validation set.

### B Background and Related Work

Methods for modeling genomic sequences can be broadly classified as alignment-free or alignment-based for functional constraints, activity predictors, and meta-predictors.

#### Alignment-free methods

Unsupervised, genome-scale language models (LMs) for eukaryotic DNA have rapidly evolved. Early examples such as DNABERT [28, 67], Nucleotide Transformer [13], HyenaDNA [47], and Caduceus [54] are all trained in a fully unsupervised manner on raw, genome-wide sequence data and have demonstrated varying strengths—some excelling at splice-site recognition, others at regulatory element prediction—depending on subtle architectural and pretraining choices [42, 38].

Building on this, foundation-scale such as Evo [46] and Evo 2 [7] have pushed both model size and context length to new extremes. Evo is a 7-billion-parameter model trained on 2.7 million prokaryotic and phage genomes (300 billion bases) with a 131 kb context window, achieving strong zero-shot performance on both predictive and generative design tasks; Evo 2 employs the StripedHyena 2 architecture and is trained autoregressively on 9.3 trillion base pairs from over 128 000 genomes spanning all domains of life, with context lengths up to 1 Mb, enabling DNA/RNA/protein multimodal prediction and de novo genome design.

Specialized alignment-free LMs focus on particular sequence classes and leverage unaligned inputs with masked language modeling (MLM) or autoregressive (GPT-style) objectives. GPN-Promoter [6] uses an MLM objective on unaligned human promoter regions to learn regulatory motifs, while SpeciesLM [19] employs MLM across 800+ unaligned genomes spanning 500 M years of evolution to capture deep conservation signals without MSAs. Other specialist models—such as plant promoter and fungal 5/3 UTR LMs [36, 19] and human 3UTR LMs [60]—also reconstruct masked nucleotides via MLM and often outperform genome-wide LMs and conservation methods like PhyloP [52] on variant effect prediction.

**Figure A1:**
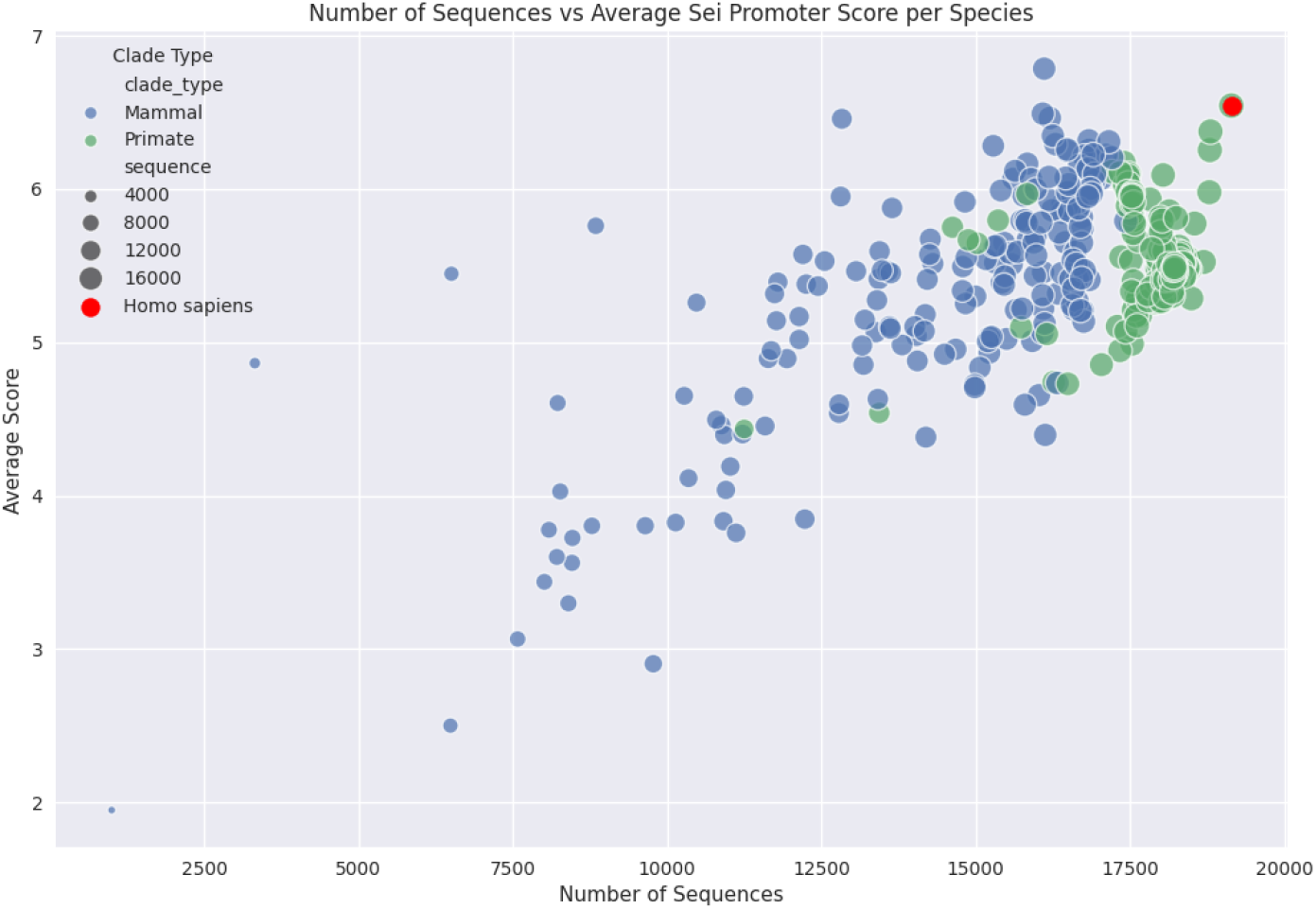
Average Promoter Sei scores plotted against the number of promoter sequences gathered for model training from the comparative genomics analysis conducted with the HAL suite. Clade types are specified by color and the red dot represents Homo sapiens. The maximum number of sequences per species is 19,254. Point sizes reflect the number of sequences.

**Figure A2:**
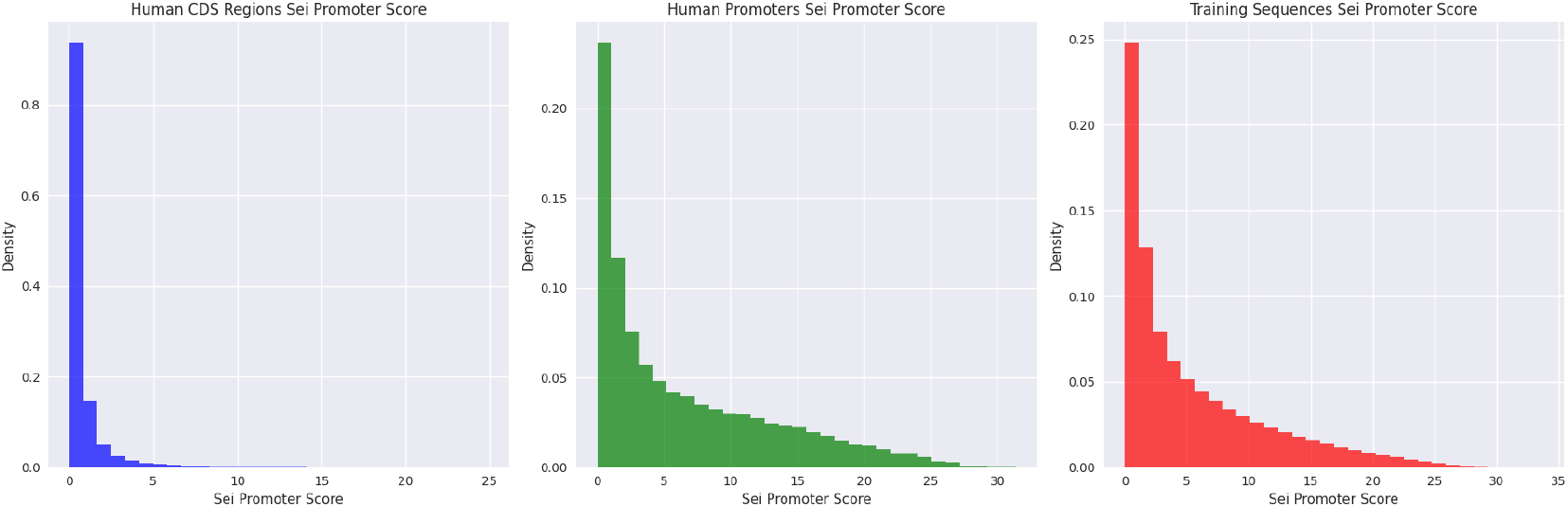
Average Promoter Sei scores were plotted for Human CDS regions, Human promoter regions, and all of the promoter data used gathered for training.

Notably, to date no autoregressive (GPT-style) models have been developed for training on mammalian promoter sequences, despite their suitability for modeling longer indels. Autoregressive protein LMs like Tranception[48] have demonstrated robust insertion and deletion effect prediction compared to MLM approaches.

#### Alignment-based methods

Multiple sequence alignments (MSAs) offer a powerful approach to understanding natural sequence variation, enabling the identification of potentially non-neutral mutations with likely functional consequences. PhyloP is an MSA-based statistical method that assigns a conservation score to each position in a sequence and compares observed substitutions to those expected under a neutral evolution model. GPN-MSA [4], a more recent development, combines whole-genome alignments with a genomic LM approach. Trained to reconstruct masked nucleotides given an MSA as input, GPN-MSA has shown improvement in SNV effect prediction compared to PhyloP. However, a major limitation of alignment-based approaches is their treatment of positions individually, which doesn’t naturally generalize to indel variants.

#### Activity Predictors & Meta Predictors

An alternative approach to unsupervised modeling of sequences involves training supervised models on measurements of sequence activity. These models often use data from high-throughput functional genomics experiments that measure various aspects of genomic function, such as expression initiation or epigenetic modifications. Models like Enformer [3] have demonstrated an understanding of factors contributing to gene expression in different cell types. However, recent studies by Sasse et al. [53] and Huang et al. [27] have shown that the performance of sequence-to-activity models such as DeepSEA [66], Basenji2 [31], and Enformer [3] in explaining expression variation between individuals due to cis-regulatory genetic variants remains limited. Another widely used method, CADD (Combined Annotation Dependent Depletion), integrates numerous genomic annotations into a single deleteriousness score[55]. However, [23] and [40] have demonstrated, comparative evaluations of meta predictors like CADD are complicated by circularity issues in their training and testing datasets leading to data leakage. As such, their performance is likely inflated due to circularity. These findings underscore the need for zero shot methods to overcome these limitations and enhance our understanding of genetic variant effects in humans.

#### B.1 Baseline details

##### B.1.1 Autoregressive models

Autoregressive LMs assign scores to sequences *s* using their log likelihood

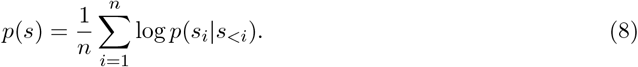

###### HyenaDNA

HyenaDNA uses base pair tokenization. For computing the cross entropy, we subset the logits and labels to the dimensions of actual nucleotides *x ∈ {A, G, C, T, N }* and exclude special tokens. We ignore the final EOS position when taking the mean over the sequence.

###### Evo1

Evo1 from [46] version *evo −* 1 *−* 131*k − base* was used. For computing the cross entropy, we subset the logits and labels to the dimensions of actual nucleotides *x ∈ {A, G, C, T, N }* and exclude special tokens. We do not apply any masking.

###### Evo2

Evo2 from [7] version *evo*2_7_*b* was used. For computing the cross entropy, we subset the logits and labels to the dimensions of actual nucleotides *x ∈ {A, G, C, T, N }* and exclude special tokens. We do not apply any masking.

##### B.1.2 Masked language models

For computational efficiency, we evaluate bidirectional masked LMs using their pseudo log likelihood,

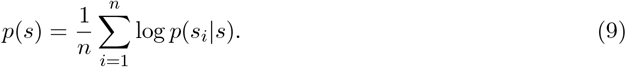

###### Caduceus

Caduceus uses base pair tokenization. For computing the cross entropy, we subset the logits and labels to the dimensions of actual nucleotides *x ∈ {A, G, C, T, N }* and exclude special tokens. We do not apply any masking.

###### Nucleotide Transformer

Nucleotide Transformer uses 6-mer tokenization. For computing the cross entropy, we subset the logits and labels to the dimensions of the 6-mer and five trailing single-base tokens and exclude special tokens. We do not apply any masking.

###### DNABERT-2

DNABERT-2 uses byte pair tokenization. For computing the cross entropy, we subset the logits and labels to the dimensions of the BPE tokens and the [UNK] token which represents *N* . Remaining special tokens are excluded. We do not apply any masking.

###### BEND – GPN

The original GPN model [5] was only trained on *Brassicales* species and is not applicable to the human genome. We instead evaluate a human GPN-based model (“Dilated ResNet”) that is included in the BEND benchmark [42]. For computing the cross entropy, we subset the logits and labels to the dimensions of actual nucleotides *x ∈ {A, G, C, T, N }*. We do not apply any masking.

###### Promoter-GPN

The original Promoter-GPN model from [6] For computing the cross entropy, we subset the logits and labels to the dimensions of actual nucleotides *x ∈ {A, G, C, T, N }*. We do not apply any masking.

###### Species-LM

Species-LM from [30] metazoa version. For computing the cross entropy, we subset the logits and labels to the dimensions of actual nucleotides *x ∈ {A, G, C, T, N }* and exclude special tokens. We do not apply any masking.

##### B.1.3 Alignment-based approaches

###### PhyloP

As they are based on an MSA, PhyloP scores are not naturally amenable to indel variants, as a change in sequence length by insertion or deletion cannot be modeled by column-wise scores. We follow gnomAD’s approach to computing PhyloP scores: For any indel, the PhyloP score of the position in the reference genome at which the indel occurs is used for the indel as a whole. Note that this inherently does not consider the actual sequence consequence of the indel - it only reflects the conservation of the position at which the indel occurs.

##### B.1.4 Activity predictors

###### Enformer

We run Enformer following the official notebook^1^. For each variant, we compute the mean difference over the sequence between the wild type and variant sequence using all human output tracks. We report the max channel to capture the largest change between the wildtype and the variant sequence.

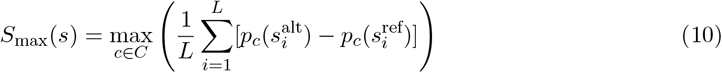

where:

*C* is the set of all human output tracks in Enformer *L* is the length of the output sequence *p*_*c*_(·) is the Enformer prediction for track 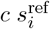 and 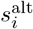 are the reference and alternate sequences at position *i*

Our Enformer evaluation computes the mean difference between wild-type and variant sequences across all human output tracks, taking the maximum as the final score. This methodology captures the maximum regulatory impact across all potential regulatory mechanisms and cell types, which is particularly important for promoter indels that may affect multiple regulatory processes simultaneously and manifest differently across diverse cellular contexts. This approach provides a more holistic assessment than the GTEx-focused SLDP regression used in the original Enformer paper, analogous to organism-scale models that consider effects across all tissue types rather than focusing on single expression outputs.

##### B.1.5 Meta Predictors

###### CADD

Combined Annotation Dependent Depletion (CADD)[34] provides a deleteriousness score across the whole genome by integrating genomic annotations and functional information, including in-silico predictions from other models. It is one of the first models to provide predictions for all single-nucleotide variants and short indels and is therefore frequently used by the community, particularly the clinical community. Of particular relevance for this work, CADD trains on population data (gnomAD frequencies), expression data (ENCODE RNAseq and epigenetic markers), transcription factor binding site annotations (ChIP transcription factor binding sites), and clinical annotations (indirectly, through training on PolyPhen2, which was itself directly trained on ClinVar labels). More information about exact features trained on can be found here: [CADD features](https://cadd.gs.washington.edu/static/ReleaseNotes_CADD_v1.7.pdf).

### C Extended Benchmark Details

#### C.1 Benchmark Implementation Details

##### C.1.1 Scoring Methodologies

To ensure fair comparisons across all models, we implement standardized scoring approaches detailed below. All models are evaluated without task-specific training or fine-tuning, though some supervised models may have been exposed to task-relevant data during their original training.

##### C.1.2 Data Leakage Assessment

While most models operate in a true zero-shot capacity, some supervised models in our evaluation have been previously exposed to task-relevant data during their training. Table A2 shows potential data leakage that occurs in supervised models for each benchmark. CADD was not used for the TFBS benchmarks due to lack of coverage.

**Table A2:**
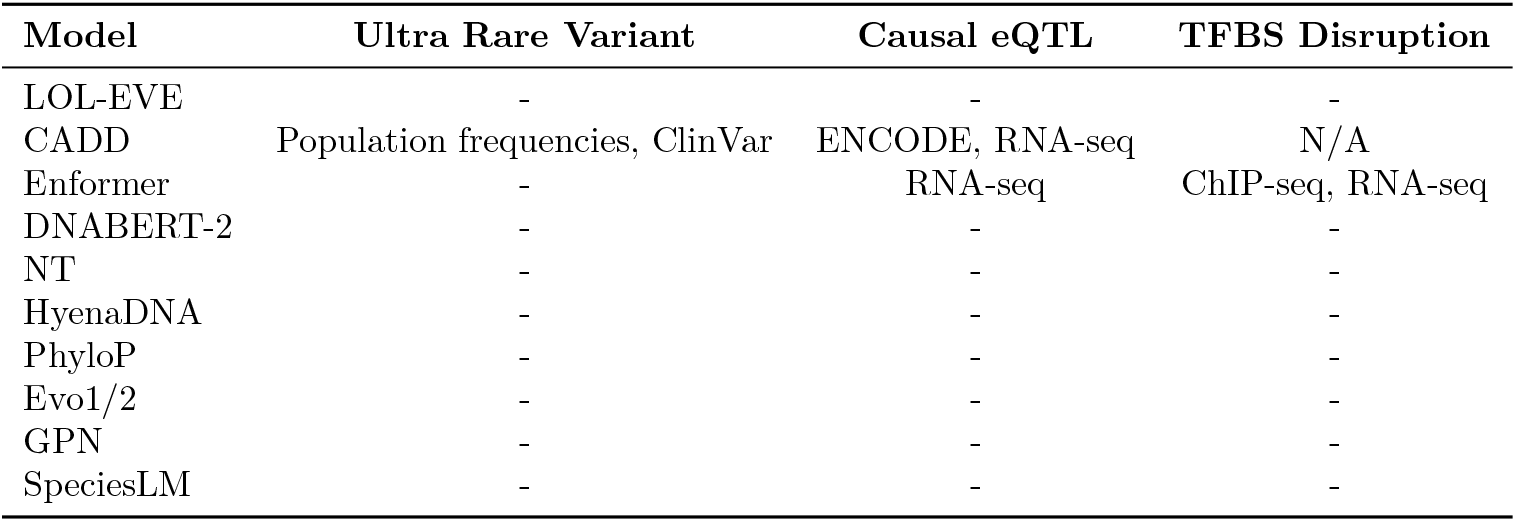
Training Data Leakage for Benchmark Tasks.

| Model | Ultra Rare Variant | Causal eQTL | TFBS Disruption |
| --- | --- | --- | --- |
| LOL-EVE | - | - | - |
| CADD | Population frequencies, ClinVar | ENCODE, RNA-seq | N/A |
| Enformer | - | RNA-seq | ChIP-seq, RNA-seq |
| DNABERT-2 | - | - | - |
| NT | - | - | - |
| HyenaDNA | - | - | - |
| PhyloP | - | - | - |
| Evo1/2 | - | - | - |
| GPN | - | - | - |
| SpeciesLM | - | - | - |

#### C.2 Ultra Rare Variant Prioritization Details

For each length category:

1. **Length bins & weights**. Partition indel lengths into 10 logarithmically spaced bins and compute the empirical bin weights *w*_*i*_.
2. **Percentiles**. For each bin *i* and percentile *p ∈ {*1, 2.5, 5, 10*}*%, compute

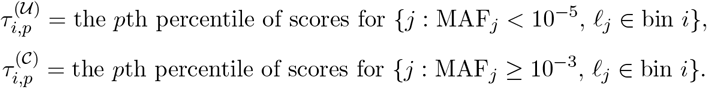
3. **Safe ratio**.

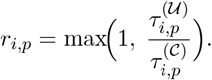
4. **Weighted mean per percentile**.

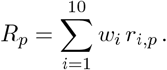
5. **Aggregate**. Report

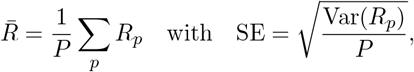

where *P* = 4 is the number of percentiles.

#### C.3 Causal eQTL Prioritization Details

##### C.3.1 Running-Mean Metric Computation

For each slippage cutoff *s*, we restrict to all indels with distance ≤ *s*. Within that subset we compute

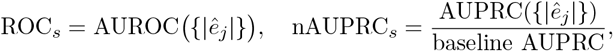

where *ê*_*j*_ is the model’s effect-score for variant *j*. Plotting ROC_*s*_ and nAUPRC_*s*_ against *s* (log-spaced) yields the cumulative performance curves in Fig. 2.

#### C.4 TFBS Disruption Detailed Methodology

##### C.4.1 Gene Stratification

We classified genes into two extreme groups using (1) evolutionary constraint—amino-acid substitution rates inferred from OrthoDB mammalian orthologs—and (2) expression variability—CV of GTEx median-TPM across tissues. “High-constraint/low-variability” genes occupy the bottom percentile in both metrics; “low-constraint/high-variability” genes occupy the top percentile. We tested robustness at 20–40% cutoffs.

##### C.4.2 TFBS Disruption Scoring

We sourced human TF motifs from JASPAR CORE [16] and retained TFs with median *TPM >* 1 in ≥ 30 GTEx tissues. Promoter sequences were scanned with PSSMs (*threshold >* 0.8) to identify binding sites; *in silico* deletions were generated, and a site was deemed “disrupted” if its post-deletion PSSM score fell below 0.8.

##### C.4.3 Balanced Comparison & Statistics

For each TF, we (a) randomly sampled equal numbers of genes from each category, (b) computed disruption scores for their TFBSs, and (c) assessed separation via point-biserial correlation. To ensure statistical reliability, we required at least 10 variants per TF per gene class for inclusion in the analysis. Only transcription factors meeting this threshold were included in the final delta accuracy calculation. P values were FDR-corrected across TFs. Finally, we report “delta accuracy” as the fraction of TFs for which high-constraint/low-variability genes scored lower (more deleterious) minus 50

#### C.5 Slippage Calculation Methodology

##### C.5.1 Rationale

DNA slippage events during replication can lead to insertions and deletions, particularly in regions with repetitive sequences or secondary structures. Understanding the relationship between model predictions and slippage propensity provides insight into whether models are learning biologically relevant mutational mechanisms versus purely statistical patterns.

##### C.5.2 Slippage Score Calculation

We implement a computational approach to estimate slippage propensity for each indel variant based on local sequence context and repetitive elements.

###### Repeat Detection Algorithm

For each variant, we extract a 20 base pair window centered on the variant position and analyze it for repetitive elements using the following approach:

1. **Homopolymer Run Detection**: We identify consecutive runs of identical nucleotides with a minimum length of 3 bases. Each homopolymer run contributes to the slippage score with a weight proportional to the square of its length.
2. **Short Tandem Repeat Detection**: We systematically search for dinucleotide, trinucleotide, and tetranucleotide repeats by:
  - Scanning the sequence with sliding windows of size 2, 3, and 4 nucleotides
  - Counting consecutive occurrences of each repeat unit
  - Requiring a minimum of 3 repeat units for classification as a tandem repeat
3. **Variant-Repeat Matching**: For each detected repeat, we check whether:
  - The deleted sequence (for deletions) matches or contains the repeat unit
  - The inserted sequence (for insertions) matches or contains the repeat unit
  - The variant position falls within the boundaries of a repeat region

###### Slippage Score Computation

The final slippage score combines contributions from all detected repeats:

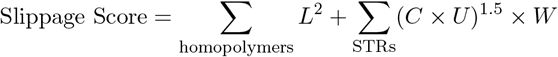

where:

- *L* = length of homopolymer run
- *C* = count of repeat units in short tandem repeat (STR)
- *U* = length of repeat unit
- *W* = weight factor: 0.8 for dinucleotides, 0.6 for trinucleotides, 0.5 for tetranucleotides

This scoring scheme gives higher weights to homopolymer runs and progressively lower weights to longer repeat units, reflecting the relative propensity for slippage in different repeat contexts.

###### Implementation Details

- Window size: 20 base pairs centered on variant position
- Minimum repeat threshold: 3 consecutive units
- Repeat unit sizes analyzed: 1-4 nucleotides
- Variants are classified as slippage-prone if they occur within or match any detected repeat region

This methodology allows us to quantitatively assess whether model predictions correlate with known mechanisms of indel formation, helping to distinguish between models that learn genuine biological constraints versus those that primarily capture mutational biases.

#### C.6 Extended Results

##### C.6.1 Ultra Rare Variant Prioritization

**Table A3:** Mean ratio and standard error for all models across indel length categories. **Bold** values indicate the best (1st place), <u>underlined</u> values indicate the second best (2nd place), and *italicized* values indicate the third best (3rd place) model performance in each category.

| Model | Small (1-2bp ) |  | Medium (3–15bp) |  | Large (16–50bp) |  |
| --- | --- | --- | --- | --- | --- | --- |
|  | Mean Ratio | Std. Error | Mean Ratio | Std. Error | Mean Ratio | Std. Error |
| CADD | *1.863* | 0.145 | *1.708* | 0.033 | **2.122** | 0.013 |
| LOL-EVE | <u>1.482</u> | 0.032 | **2.125** | 0.051 | <u>2.097</u> | 0.109 |
| GPN-Promoter | **2.297** | 0.025 | <u>1.885</u> | 0.032 | 1.490 | 0.073 |
| speciesLM | 1.220 | 0.017 | 0.986 | 0.076 | *1.740* | 0.030 |
| HyenaDNA-tiny | 1.323 | 0.028 | 1.410 | 0.009 | 1.368 | 0.041 |
| HyenaDNA-small | 1.283 | 0.016 | 1.350 | 0.007 | 1.295 | 0.032 |
| HyenaDNA-medium-160k | 1.373 | 0.011 | 1.406 | 0.018 | 1.260 | 0.027 |
| HyenaDNA-medium-450k | 1.335 | 0.025 | 1.348 | 0.023 | 1.308 | 0.038 |
| HyenaDNA-large | 1.345 | 0.018 | 1.340 | 0.017 | 1.207 | 0.008 |
| GPN | 1.198 | 0.020 | 1.299 | 0.079 | 1.269 | 0.044 |
| NT-v2-500m | 1.131 | 0.024 | 1.209 | 0.041 | 1.284 | 0.014 |
| NT-500m | 1.059 | 0.021 | 1.155 | 0.005 | 1.275 | 0.039 |
| NT-2.5b-multi | 1.022 | 0.014 | 1.068 | 0.015 | 1.330 | 0.041 |
| NT-2.5b-1000g | 1.013 | 0.017 | 1.091 | 0.031 | 1.247 | 0.075 |
| Caduceus-ps | 0.943 | 0.018 | 1.079 | 0.049 | 1.310 | 0.125 |
| Caduceus-ph | 1.028 | 0.045 | 1.119 | 0.071 | 1.317 | 0.161 |
| DNABERT-2 | 1.069 | 0.012 | 0.975 | 0.017 | 1.067 | 0.007 |
| PhyloP | 1.067 | 0.037 | 1.044 | 0.013 | *1.344* | 0.027 |
| Evo1 | *1.262* | 0.033 | 1.336 | 0.102 | 1.060 | 0.034 |
| Evo2 | 0.822 | 0.041 | 0.777 | 0.015 | 1.007 | 0.011 |
| Enformer | 1.000 | 0.000 | 1.000 | 0.000 | 1.000 | 0.000 |
| GC Content $\Delta$ | 1.005 | 0.001 | 1.027 | 0.010 | 0.876 | 0.007 |
| Distance TSS | 1.000 | 0.000 | 0.921 | 0.008 | 1.226 | 0.329 |

**Table A4:**
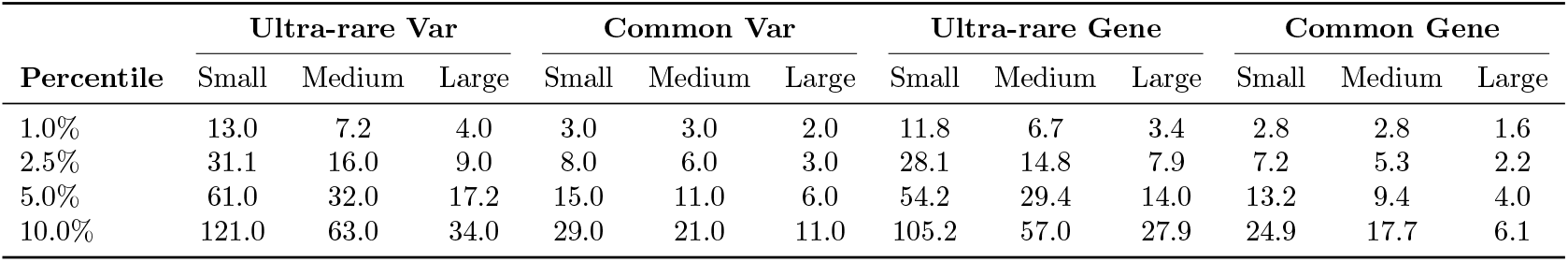
Average counts per model of variants (Var) and genes (Gene), stratified by rarity (ultra-rare vs common), percentile, and indel size. No model reported zero counts in any of these splits.

| Percentile | Ultra-rare Var |  |  | Common Var |  |  | Ultra-rare Gene |  |  | Common Gene |  |  |
| --- | --- | --- | --- | --- | --- | --- | --- | --- | --- | --- | --- | --- |
|  | Small | Medium | Large | Small | Medium | Large | Small | Medium | Large | Small | Medium | Large |
| 1.0% | 13.0 | 7.2 | 4.0 | 3.0 | 3.0 | 2.0 | 11.8 | 6.7 | 3.4 | 2.8 | 2.8 | 1.6 |
| 2.5% | 31.1 | 16.0 | 9.0 | 8.0 | 6.0 | 3.0 | 28.1 | 14.8 | 7.9 | 7.2 | 5.3 | 2.2 |
| 5.0% | 61.0 | 32.0 | 17.2 | 15.0 | 11.0 | 6.0 | 54.2 | 29.4 | 14.0 | 13.2 | 9.4 | 4.0 |
| 10.0% | 121.0 | 63.0 | 34.0 | 29.0 | 21.0 | 11.0 | 105.2 | 57.0 | 27.9 | 24.9 | 17.7 | 6.1 |

##### C.6.2 Causal eQTL Prioritization

**Figure A3:**
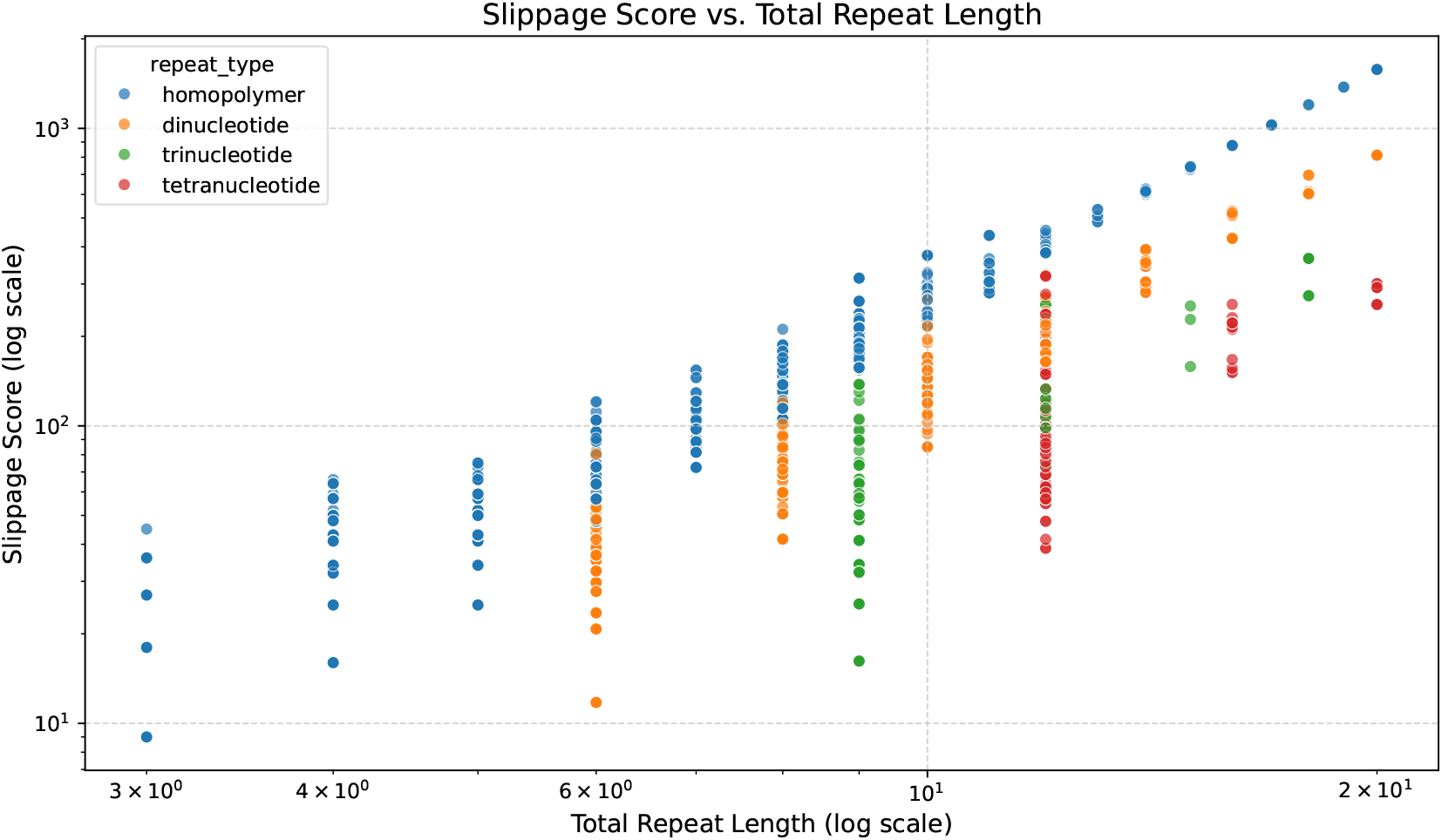
The slippage score assigned to different repeat types.

**Table A5:** Full breakdown of gene and variant counts per pip threshold and slippage threshold.

| PIP Threshold | Slippage Threshold |  |  |  |  |  |  |  |  |  |  |  |
| --- | --- | --- | --- | --- | --- | --- | --- | --- | --- | --- | --- | --- |
|  | 10 | 20 | 30 | 40 | 50 | 75 | 100 | 200 | 300 | 400 | 500 | inf |
| *Background Variant Counts* |  |  |  |  |  |  |  |  |  |  |  |  |
| 0.001 | 9 | 20 | 22 | 24 | 25 | 32 | 35 | 47 | 48 | 50 | 51 | 52 |
| 0.01 | 315 | 506 | 654 | 774 | 883 | 1036 | 1135 | 1796 | 1860 | 1890 | 1900 | 1933 |
| 0.05 | 827 | 1306 | 1681 | 1951 | 2236 | 2613 | 2834 | 4397 | 4560 | 4626 | 4649 | 4719 |
| *Causal Variant Counts* |  |  |  |  |  |  |  |  |  |  |  |  |
| 0.001 | 12 | 19 | 31 | 34 | 38 | 48 | 49 | 76 | 81 | 82 | 82 | 82 |
| 0.01 | 23 | 32 | 50 | 60 | 72 | 89 | 94 | 140 | 147 | 148 | 148 | 148 |
| 0.05 | 35 | 50 | 77 | 95 | 115 | 144 | 157 | 224 | 240 | 243 | 243 | 243 |
| *Background Gene Counts* |  |  |  |  |  |  |  |  |  |  |  |  |
| 0.001 | 9 | 19 | 21 | 23 | 24 | 31 | 34 | 45 | 46 | 48 | 49 | 50 |
| 0.01 | 298 | 472 | 601 | 701 | 794 | 917 | 994 | 1533 | 1570 | 1591 | 1594 | 1608 |
| 0.05 | 766 | 1189 | 1501 | 1712 | 1924 | 2211 | 2370 | 3513 | 3591 | 3624 | 3629 | 3650 |
| *Causal Gene Counts* |  |  |  |  |  |  |  |  |  |  |  |  |
| 0.001 | 12 | 19 | 31 | 34 | 38 | 48 | 49 | 76 | 81 | 82 | 82 | 82 |
| 0.01 | 22 | 31 | 49 | 58 | 70 | 87 | 92 | 137 | 144 | 145 | 145 | 145 |
| 0.05 | 34 | 49 | 76 | 93 | 112 | 139 | 151 | 215 | 229 | 232 | 232 | 232 |

**Figure A4:**
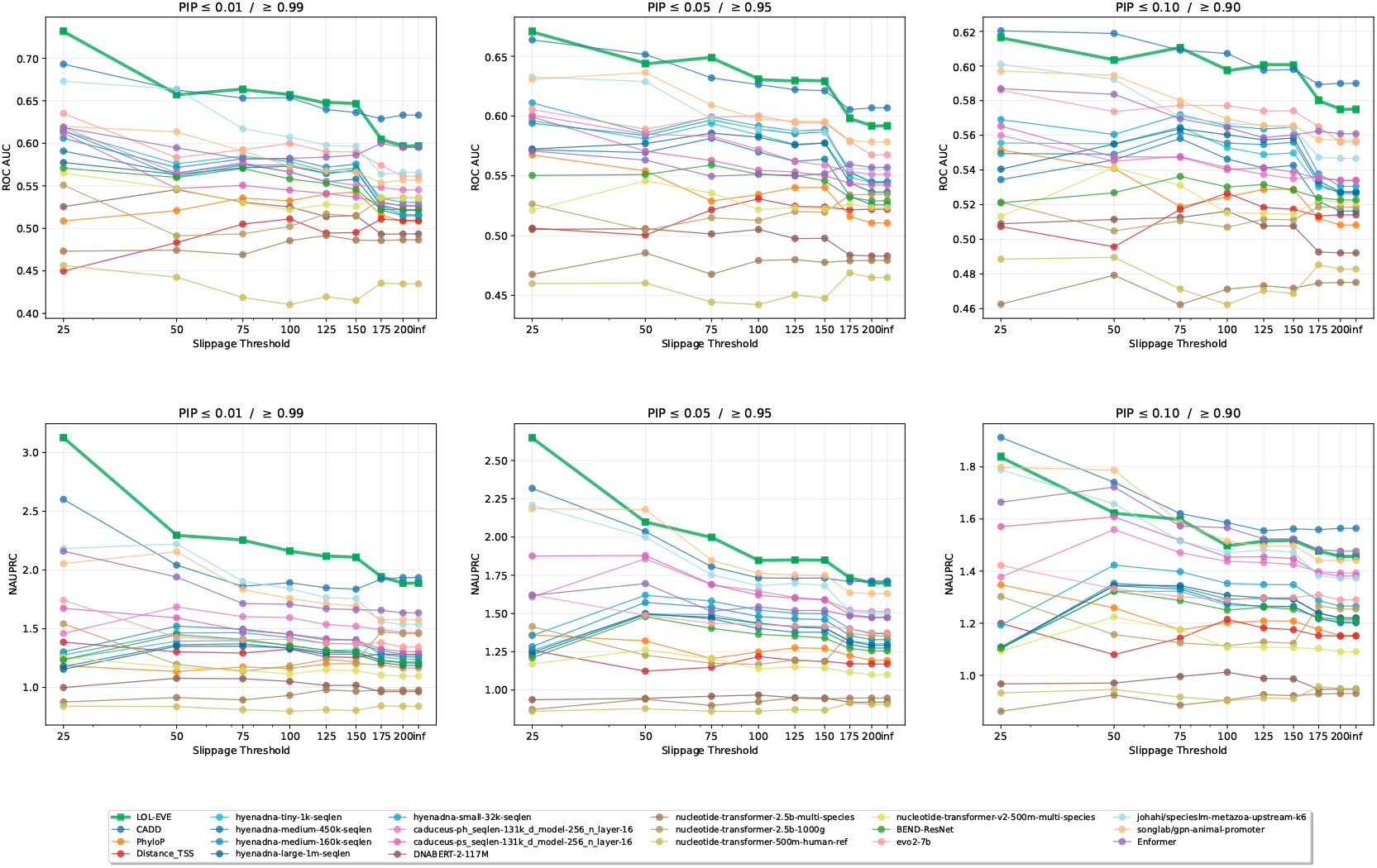
Cumulative causal-eQTL performance curves (running-mean AUROC and normalized AUPRC) as a function of slippage cutoff (log scale).

**Figure A5:**
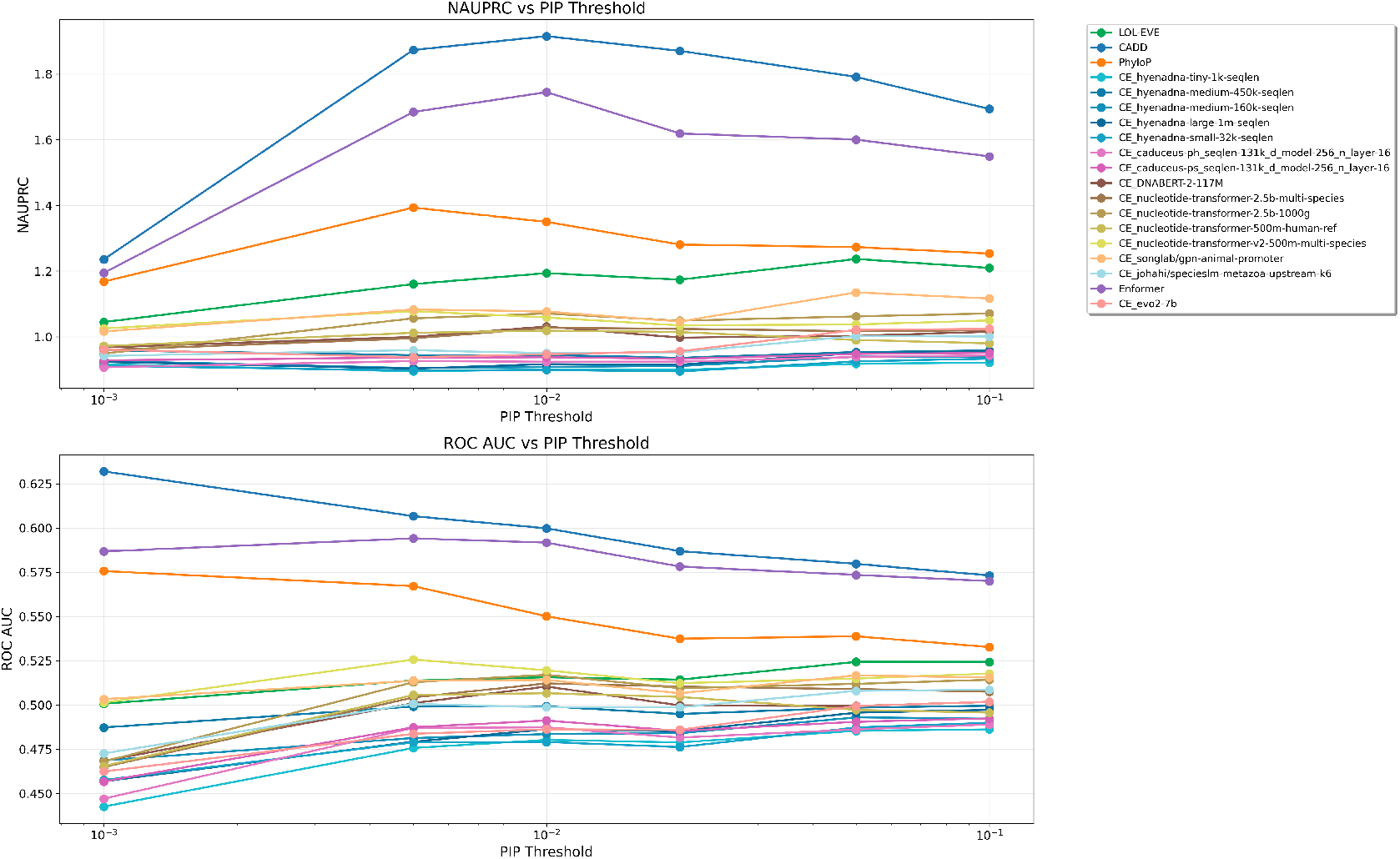
SNP prioritization: Normalized AUPRC, and ROC AUC for each model. Full breakdown of variants per cutoff are in Table A6.

**Table A6:** SNP variant class counts across different PIP thresholds for variant classification.

| PIP Threshold | Total | Causal | Background | Class Ratio |
| --- | --- | --- | --- | --- |
| 0.001 | 758 | 444 | 314 | 1.41 |
| 0.005 | 6,200 | 619 | 5,581 | 9.02 |
| 0.01 | 12,708 | 733 | 11,975 | 16.34 |
| 0.02 | 20,577 | 893 | 19,684 | 22.05 |
| 0.05 | 30,271 | 1,213 | 29,058 | 23.96 |
| 0.1 | 35,597 | 1,403 | 34,194 | 24.37 |

##### C.6.3 TFBS Disruption

**Figure A6:**
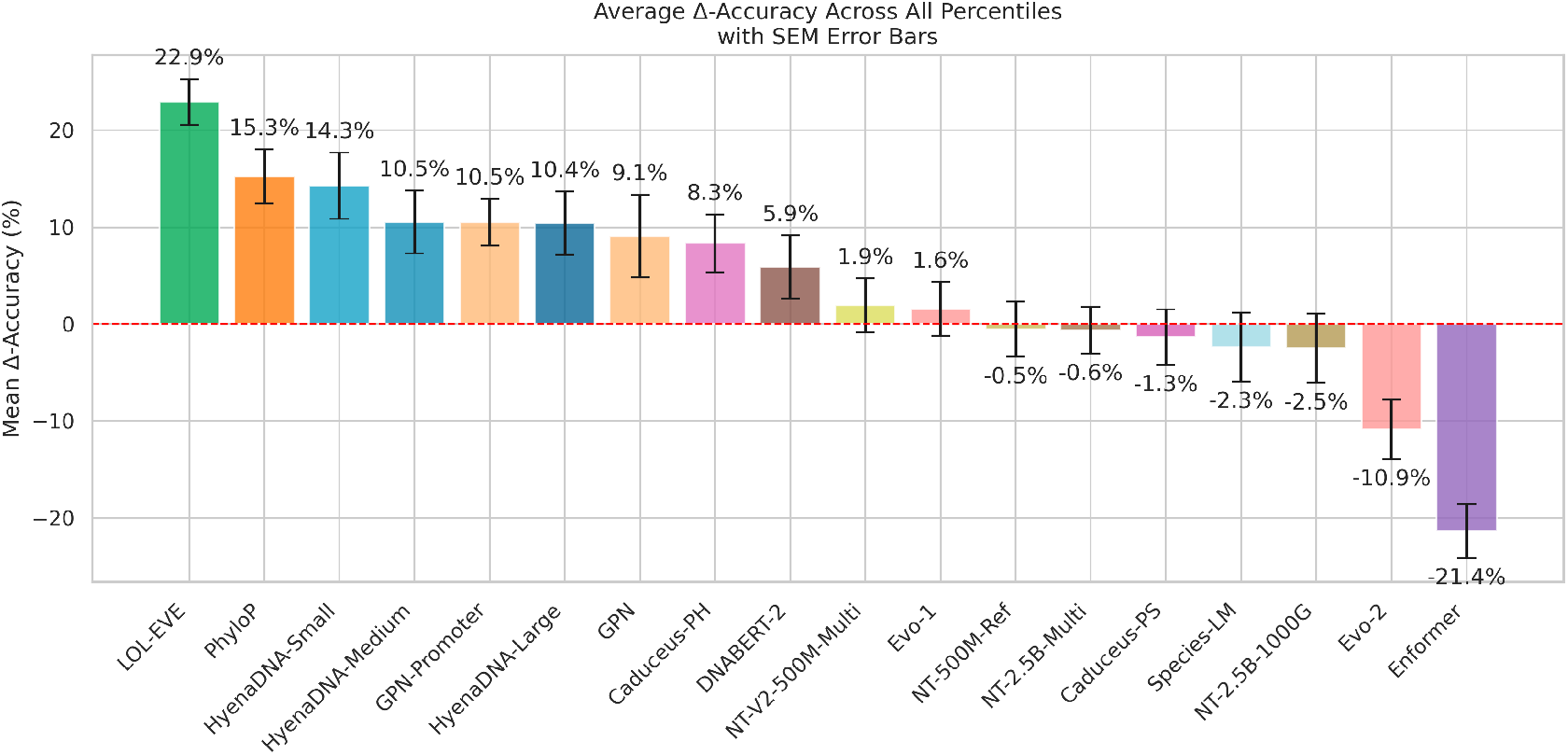
All models show for TFBS task.

**Figure A7:**
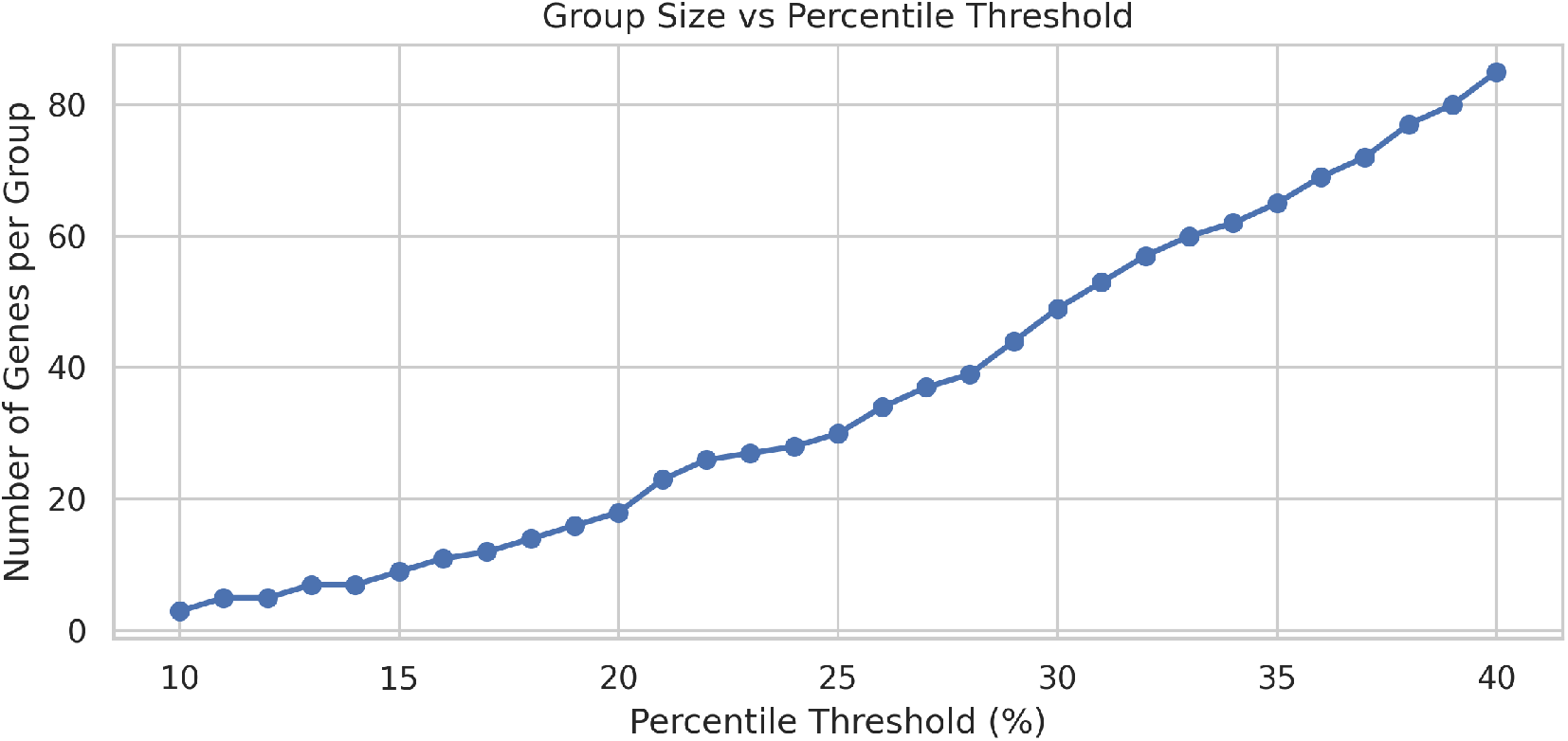
Cumulative gains of Genes per percentile threshold.

**Figure A8:**
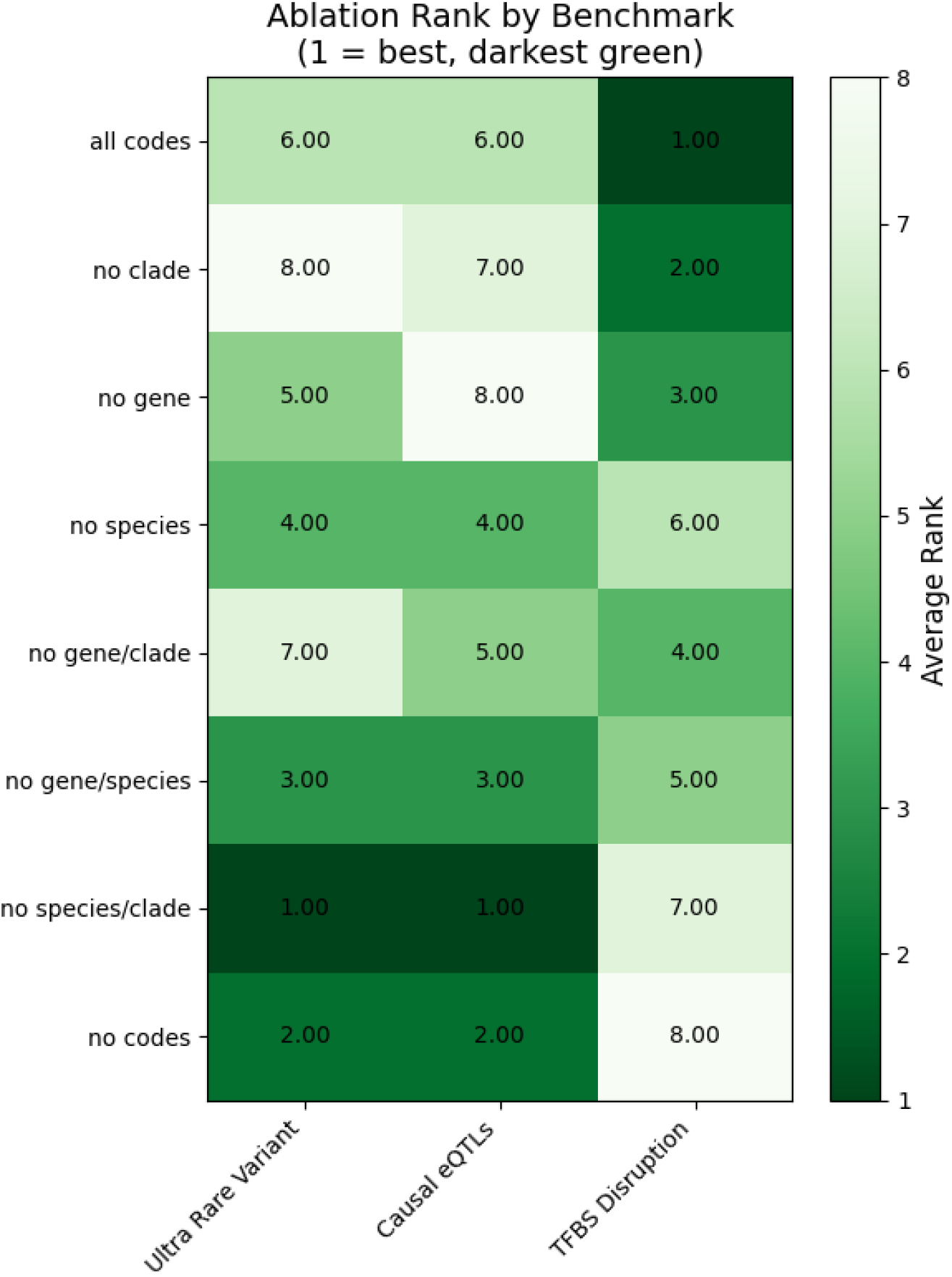
ABlation of control codes in LOLEVE for the different Benchmarks. Pip Thresholds were averaged for the Causal eQTL task, and indel sizes ranges were averaged for the Ultra Rare Variant Prioritization Task.

### D Genomics England Data Processing

#### D.1 Variant Filtering and Promoter Annotation

Variants from trio VCF files were intersected with promoter regions using bedtools intersect. To handle variants that overlap multiple promoters, each variant-promoter combination was treated as a separate entry in the filtered VCF files. This approach ensures that variants falling within overlapping promoter regions are scored independently in each genomic context, preserving the biological relevance of promoter-specific effects. Each variant entry was annotated with the corresponding promoter name and strand information in the VCF INFO field.

#### D.2 Haplotype Construction for Language Model Scoring

For each trio, we constructed haplotypes by combining variants according to their parental inheritance patterns. Variants were classified as maternally inherited, paternally inherited, or de novo mutations (DNMs) based on trio genotype analysis. For inherited variants, haplotypes were built by grouping variants from the same parental origin within each promoter region. For DNMs, we generated all possible haplotype combinations by testing the de novo variant in combination with both maternally and paternally inherited background variants within the same promoter. This exhaustive approach allows assessment of how DNMs interact with different inherited variant backgrounds when scored by genomic language models. Haplotypes were scored using LOL-EVE, HyenaDNA, and Evo2.

#### D.3 Stat Analysis

To test for enrichment of low-scoring haplotypes in known developmental disorder genes, we calculated odds ratios across different score thresholds. First, we selected the most deleterious scoring haplotype per individual. Then, for each threshold, we calculated how many people had a promoter below or above that threshold in front of a known developmental disorder gene according to the DDG2P [15]. We then compared the ratio of these two to the number of people who had their worst scoring promoter in front of a non-developmental disorder gene below and above the threshold.

**Figure A9:**
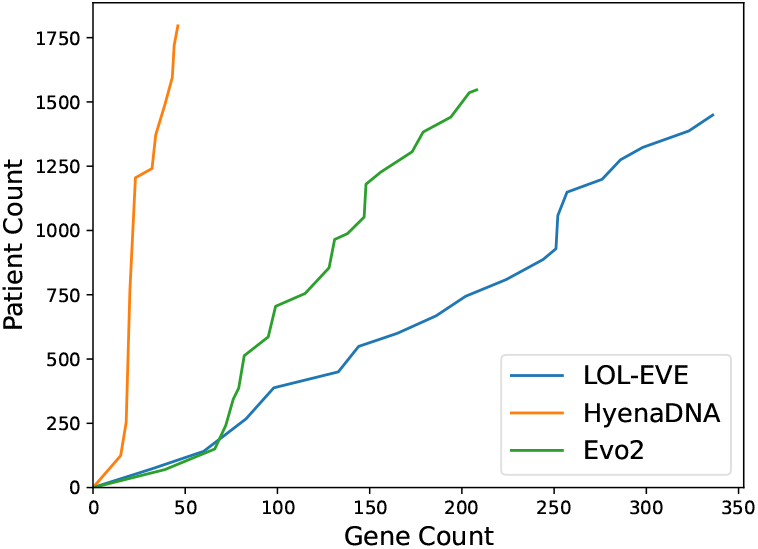
Across LLR percentile thresholds described in subsection D.3 the Gene Count and Patient Count are reported across LOL-EVE, HyenaDNA, and Evo2.

**Figure A10:**
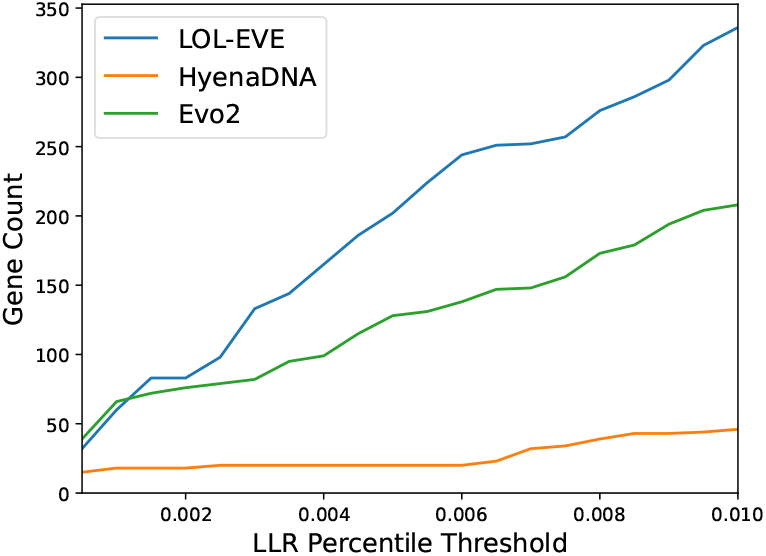
Across LLR percentile thresholds described in subsection D.3 the Gene Count is reported across LOL-EVE, HyenaDNA, and Evo2.

## Footnotes

1 https://github.com/google-deepmind/deepmind-research/blob/master/enformer/enformer-usage.ipynb

